# Attenuated foreign body response to subcutaneous implant in regenerative spiny mice (*Acomys*)

**DOI:** 10.1101/2022.08.12.503776

**Authors:** Janak Gaire, Michele Dill, Valentina Supper, Chelsey S. Simmons

## Abstract

Spiny mice (*Acomys*) can regenerate after injury with minimal fibrosis. Whether *Acomys* retains the fibrosis-free feature in response to implanted devices is unknown, so we implanted polydimethylsiloxane (PDMS) subcutaneously in *Acomys* and *Mus*, a non-regenerative counterpart. In *Acomys*, we found reduced myeloid cell infiltration, fibroblast activation, and collagen deposition around the PDMS implant. These results suggest that *Acomys* can regulate FBR and may hold the key to improving implant lifetime and functionality.

## Introduction

Most adult mammals, including humans, cannot regenerate damaged organs and tissues. The functionality of damaged or malfunctioning organs can be augmented, to some extent, by using implantable technologies such as insulin pumps and glucose sensors to manage diabetes^1^, artificial pacemakers for cardiovascular issues^2^, and neural implants to restore lost sensory and motor functions^3^. While the aforementioned implantable devices (and many others) can significantly improve patient quality of life, they do not function reliably long-term. This decline in functionality is attributed to the foreign body response (FBR), a complex sequence of biochemical signaling that is initiated immediately after implantation and persists for the lifetime of the device. FBR is characterized by the recruitment and activation of immune cells (e.g., neutrophils and macrophages) and stromal cells (e.g., fibroblasts), which results in fibrotic encapsulation around the implanted device and often leads to complete device failure.

Intervention strategies such as modification of implant surface topography and stiffness or the use of anti-inflammatories^4–6^ can reduce the severity of FBR; however, these modifications still do not allow seamless integration of biomedical implants into the host tissue. Leveraging mechanisms underlying scar-free regeneration in regenerative organisms such as Spiny mice (*Acomys*) may lead us to identify new solutions to modify the body’s response to biomedical implants. Unlike most mammalian adults, *Acomys* fully restores tissue architecture and functionality after a wide range of injuries by evading fibrosis^7–9^.

While the biological underpinnings of scar-free regeneration in *Acomys* are yet to be elucidated, previous studies suggest differential activation of immune cells, fibroblasts, and collagen remodeling in the *Acomys* wound bed play key roles. Specifically, the regenerating *Acomys* wound bed contains fewer immune cells like neutrophils^10,11^ and pro-inflammatory macrophages^8,10^, as well as fewer myofibroblasts^7,12^ and decreased collagen deposition^10^. Furthermore, *Acomys* fibroblasts cultured on stiffer substrates do not display a myofibroblast phenotype^13^ nor do they differentiate into myofibroblasts when treated with transforming growth factor beta1 (TGF-b1)^12^, suggesting a resistance to myofibroblast differentiation even in the presence of pro-fibrotic stimuli. These findings in *Acomys*, along with the similarities between fibrosis- and FBR-associated pathways, suggest that *Acomys* may demonstrate a reduced FBR to implanted biomedical devices. However, to date, the response of *Acomys* to biomedical implants remains unknown.

To investigate the response of *Acomys* to implanted biomaterials, we fabricated 2×2-mm implants using polydimethylsiloxane (PDMS), a polymer commonly used in commercially available, FDA-approved devices^5,14^. We implanted PDMS sheets under the dorsolateral skin of *Acomys* and *Mus*, a non-regenerative, scar-forming counterpart. After 5 and 30 days post-implantation (DPI), the implants and surrounding tissue were excised and assessed via histology to compare changes during the inflammatory and tissue-remodeling stages.

## Results

### Acomys had fewer myeloid cells and fibroblasts surrounding the implant at 5 and 30 DPI and reduced cellularity at 30 DPI

To assess cellularity surrounding the PDMS implant, tissue sections were stained with H&E stain. Compared to a distant site, elevated levels of total nuclei count were present near the implant across all groups (**Figure 1B-C**). At 5 DPI, the fold-change of total nuclei, calculated by normalizing the total cell count surrounding the implant to the cell count away from the implant site, were comparable between *Mus* and *Acomys* (2.86 ± 0.55 versus 2.58 ± 0.45, p = 1.0). However, at 30 DPI, the total nuclei remained elevated in *Mus* but decreased in *Acomys* (3.31 ± 0.46 versus 1.77 ± 0.16, p = 0.02).

**Figure 1.**
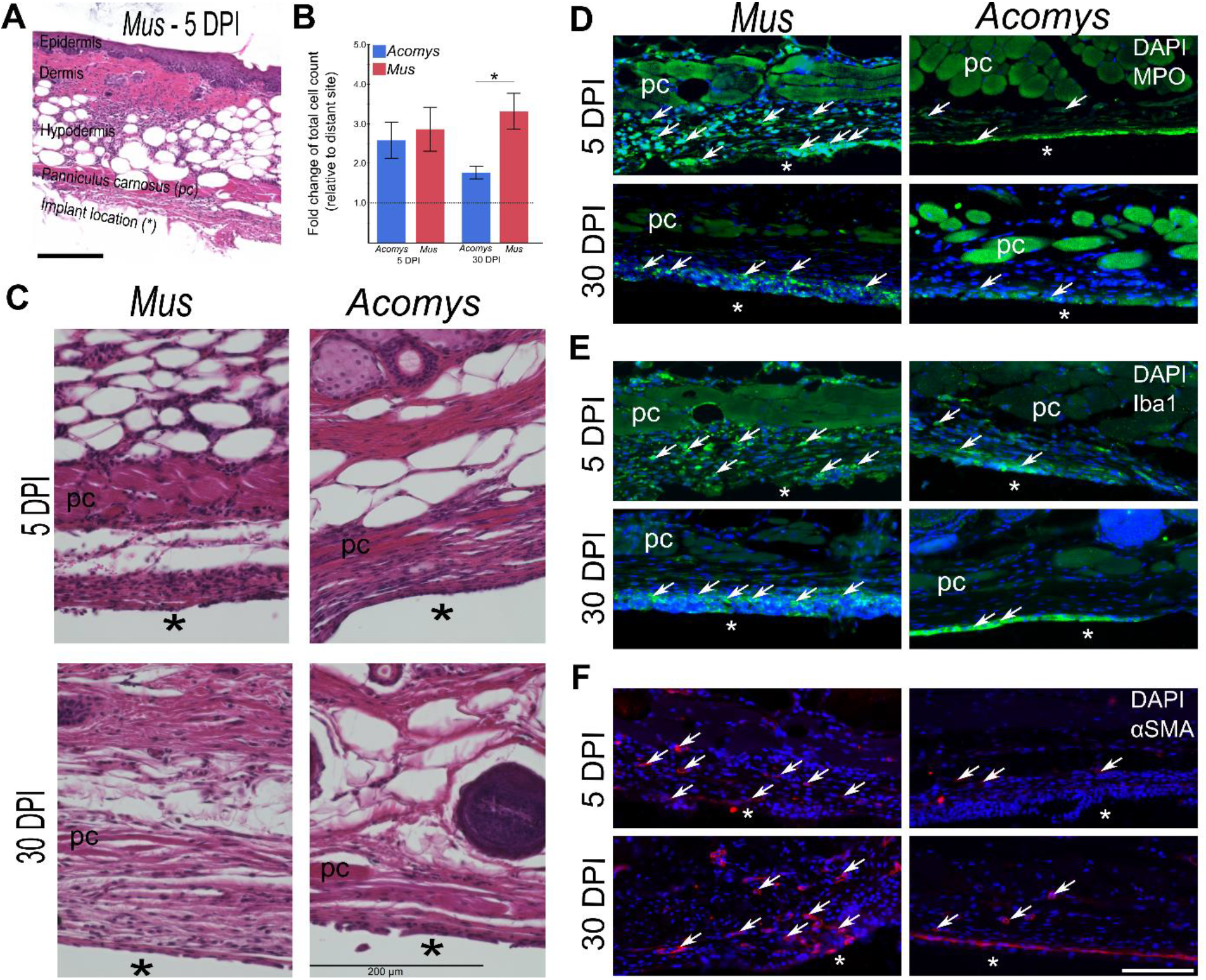
*Acomys* display reduced cellularity, fewer immune cells, and fewer activated myofibroblasts surrounding the PDMS implant. **A.** An H&E-stained image of *Mus* skin section (5 DPI) showing three different layers of skin, panniculus carnosus (pc) – a muscle layer below the hypodermis, and implant location (*). Scale bar = 200 μm. **B.** Fold change of total cell count was calculated by obtaining the average nuclei count surrounding the implant at 5-7 different locations per section (2-3 sections per animal) normalized to average cell count in a similar location away from the implant site. Differences among groups was analyzed using non-parametric Kruskal-Wallis test followed by multiple comparisons using the Steel-Dwass method. Error bars in B represent the standard error of the mean. * p = 0.02. **C.** Representative images of *Mus* and *Acomys* skin sections from 5 and 30 days post-implant (DPI) stained with H&E. **(D-F)** Representative images of *Mus* and *Acomys* skin sections from 5 and 30 DPI immunolabeled with **(D)** Myeloperoxidase, a marker for myeloid cells including neutrophils, (**E**) Iba1, a marker for macrophages, and (**F**) alpha smooth muscle actin (αSMA), a marker for activated fibroblasts. Arrows in the figures represent example cells positive for each marker. Asterisk (*) denotes implant location. Scale bars in C & F are 200 μm & F = 100 μm, respectively.

To further explore the type of cell populations near the implant site, tissue sections were immunolabeled using markers for myeloid cells (myeloperoxidase, MPO; ionized calcium-binding adapter molecule 1, Iba1) and activated fibroblasts (alpha smooth muscle actin, αSMA). We observed substantial differences in the distribution of immune cells and fibroblasts between *Mus* and *Acomys*. In *Mus*, many nucleated cells surrounding the implant were myeloid cells including neutrophils and macrophages as indicated by the abundant MPO+ (**Figure 1D**) and Iba1+ (**Figure 1E**) cells, respectively, at 5 DPI. These myeloid cell populations persisted through 30 DPI, albeit in reduced quantities that were largely localized at the implant vicinity. In contrast, very few MPO+ and Iba1+ cells were present in *Acomys* surrounding the implant at either time point. Furthermore, after immunolabeling the skin sections for αSMA, a marker for myofibroblasts, many αSMA+ cells were present in *Mus* at both 5 and 30 DPI whereas in *Acomys*, there were very few αSMA+ cells that were mostly confined to blood vessels (**Figure 1F**).

### Reduced fibrotic encapsulation was observed in Acomys at 30 DPI

Next, the extent of fibrotic encapsulation was characterized by staining skin sections with Masson’s trichrome stain, a dye specific for collagen. Qualitatively, at 5 DPI, we observed a relatively weak and diffuse collagen signal peri-implant in both species (**Figure 2A**). At 30 days, excessive deposition of extracellular matrix was observed in *Mus*, as indicated by denser staining and the presence of thick bundles of collagen fibers fully encapsulating the PDMS implant. In contrast, at 30 DPI, collagen signal surrounding the implant in *Acomys* appear more or less similar to 5 DPI. *Acomys* samples did not display the dense, thick bundles of collagen surrounding the implant seen in *Mus* (**Figure 2A**). To quantify fibrotic encapsulation, we measured the distance between the edge of the implant on the superficial side (facing the panniculus carnosus, PC) and the bottom of PC as outlined in **Figure 2B** at 5 different locations. At 5 DPI, the collagen thickness was comparable between the two species (p = 0.84).

**Figure 2.**
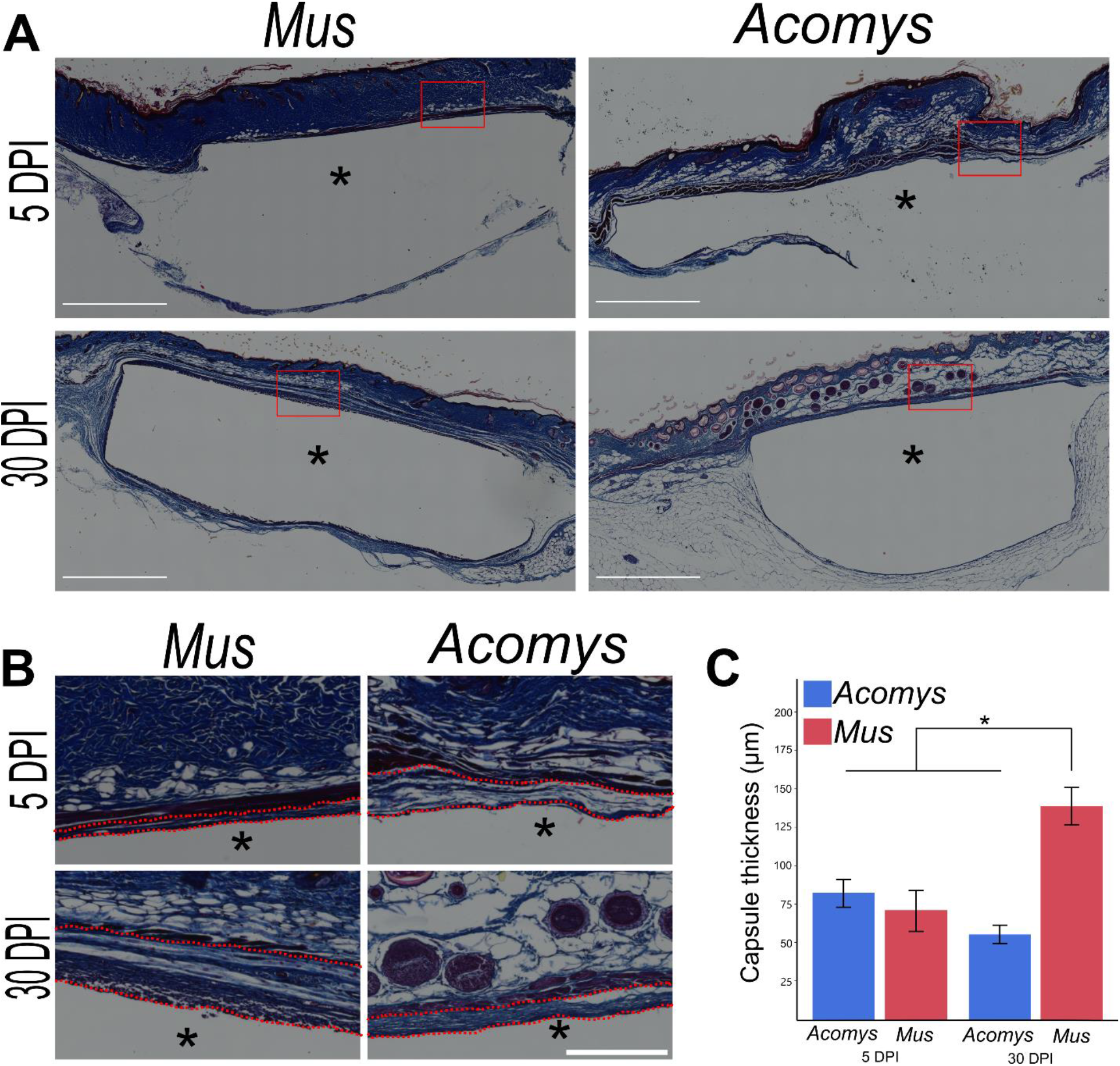
Reduced fibrotic encapsulation in *Acomys* at 30 days post-implant (DPI). **A.** Representative stitched tile-scanned images of *Mus* and *Acomys* skin sections from 5 and 30 DPI stained with Masson’s Trichrome to visualize collagen. Asterisk (*) denotes implant location. Scale bars = 500 μm. **B.** Magnified images from red boxes in A. Red dashed lines drawn at edge of the implant (on superficial side) facing panniculus carnosus layer and the bottom of panniculus carnosus layer represent location of capsule thickness measurement. Scale bar = 100 μm. **C.** Quantification of capsule thickness. Differences among groups was analyzed using non-parametric Kruskal-Wallis test followed by multiple comparisons using with the Steel-Dwass method. Error bars represent the standard error of the mean. *p<0.001.

However, at 30 DPI, *Mus* had significantly greater collagen thickness compared to *Acomys* (p < 0.0001, **Figure 2B and 2C**). In *Mus*, collagen thickness was also significantly higher at 30 DPI compared to 5 DPI (p = 0.02) whereas no statistically significant differences were observed between 5 and 30 DPI in Acomys (p = 0.17).

## Discussion

Here, we report for the first time, that *Acomys* extends its “fibrosis-free” regenerative response to implanted devices. We placed a PDMS implant subcutaneously in *Acomys* and compared histological changes surrounding the implant with *Mus*, a non-regenerative, scar-forming counterpart. Compared to *Mus*, we found minimal activity of immune cells and fibroblasts and reduced collagen deposition surrounding the PDMS implant in *Acomys*.

Similar to the blunted immune response seen after injury^9^, *Acomys* display a dampened immune response in the presence of implanted devices. While there were no differences in the total nuclei count initially between *Mus* and *Acomys*, we found substantial differences in cell composition surrounding the PDMS implant during the initial inflammatory stage (5 DPI). Specifically, we observed reduced inflammatory activity at 5 DPI in *Acomys* as indicated by the presence of few MPO+ myeloid cells and Iba1+ macrophages around the implant. This could have led to reduced fibrotic encapsulation as macrophages are known to be involved in fibroblast activation during FBR^15^. Moreover, during the tissue remodeling phase (30 DPI), the overall nuclei count was lower in *Acomys*, and immune cells were near absent at the peri-implant region, indicating the absence of persistent inflammatory response even in the presence of a foreign object. In *Mus*, however, implant-induced chronic inflammation was observed as indicated by increased total cell count and the continued presence of MPO+ and Iba1+ cells surrounding the implant. Studying altered immune cell recruitment and activation in *Acomys* will continue to be necessary in improving our understanding of regeneration and may be extended to the design of biomaterials to prevent FBR.

While the collagen levels were similar between *Mus* and *Acomys* at 5 DPI, there was a significantly higher amount of collagen deposited near the implant in *Mus* at 30 DPI. Furthermore, in *Acomys*, collagen content surrounding the implant was similar between 5 and 30 DPI. Reduced collagen deposition around the PDMS implant in *Acomys* is likely attributed to reduced myofibroblast activation as there were very few αSMA positive cells at both 5 and 30 DPI. The lack of myofibroblast activation in *Acomys* could be due to either reduced mechanical activation, reduced immune cell signaling, or a combination of both. Stewart *et al*. previously demonstrated that *Acomys* fibroblasts do not differentiate into myofibroblasts in response to increasing substrate stiffness^13^. Furthermore, softer implants are also known to result in reduced myofibroblast activation and reduced fibrotic encapsulation^6^. It is possible that mechanotransduction may be impeded in *Acomys* fibroblasts, causing cells to behave as though they are in contact with soft substrates and preventing their activation and differentiation into myofibroblasts. Therefore, uncovering the mechanisms behind impeded mechanotransduction in *Acomys* fibroblasts may allow us to translate the findings to prevent FBR in scar-forming organisms.

Overall, *Acomys* display reduced foreign body response to PDMS implants, as indicated by fewer MPO+, Iba1+, and αSMA+ cells and reduced collagen deposition surrounding the PDMS implant. These observations are similar to those reported in other *Acomys* injury models, implicating the potential involvement of similar pathways in scar-free regeneration and FBR. Future studies geared towards identifying a mechanistic basis for scarless healing in *Acomys* could prove beneficial in designing biomedical implants that reduce FBR, thereby increasing their efficacy and functional longevity.

## Materials and Methods

### 2.1 Animals

All animal experiments were conducted in accordance with United States Department of Agriculture (USDA) and National institute of Health (NIH) guidelines and were approved by the Institutional Animal Care and Usage Committee (IACUC) at the University of Florida. Spiny mice (*Acomys cahirinus*) were procured from a breeding colony at the University of Florida and CD-1 laboratory mice (*Mus musculus*) were purchased from Charles River Laboratories (Wilmington, MA). Animals of both sexes were included in this study.

### 2.2 PDMS implant preparation

PDMS implants were prepared by mixing two components of the Sylgard™ 182 silicone elastomer kit (Dow Corning) in a 10:1 ratio by weight. The mixture was poured into a petri dish to obtain approximately 1 mm thickness, degassed for an hour, and cured overnight at 70°C. Implants were then cut into approximately 2X2 mm squares, rinsed with Milli-Q water to remove any debris, and autoclaved for sterilization. Once sterilized, the implants were kept in sterile 0.9% sodium chloride solution (SteriCare™ Solutions) at room temperature.

### 2.3 Surgical procedures

Animals were anesthetized using isoflurane (initial induction at 3% isoflurane in O2 and maintenance at 1-2% isoflurane in O2 during surgery). The surgical site was shaved using an electric razor and cleaned using three alternating washes of chlorohexidine and saline. An approximately 2 cm incision was made on the dorsolateral side and a subcutaneous pocket was created via blunt dissection using sterile surgical scissors. A PDMS implant, prepared as described in 2.2, was placed into the pocket and the incision site was sealed using a 4-0 non-absorbable polyamide surgical suture (Redilon ^®^ Pro, Myco Medical) and a few drops of surgical glue (3M Vetbond™ Tissue adhesive). Once ambulatory, animals were returned to their respective cages. Meloxicam (20 mg/kg) was administered subcutaneously as an analgesic until 48 hours after the surgery.

### 2.4 Histological procedures (tissue processing, staining, and immunohistochemistry)

At 5 days (N=3 *Mus* and *Acomys*) or 30 days (N=4 for *Acomys* and N = 5 for *Mus*) post-implantation, animals were euthanized by CO2 asphyxiation and decapitated. Skin was excised to carefully retrieve implants along with the surrounding tissues and prepared for standard histological preparations. In brief, samples were fixed in 4% paraformaldehyde (PFA) for 24 hours followed by three washes with phosphate buffer solution (PBS) to remove excess PFA. Tissue samples containing implants were dehydrated by incubating in a series of graded alcohol (70%, 95%, and 100%), xylene, and paraffin wax. Samples were then cut in halves (approximately from the center) and PDMS implants were carefully removed prior to embedding in paraffin blocks. Skin samples embedded in paraffin blocks were sectioned into 5-10 μm thick sections using a microtome (Microm HM355S, Thermo Scientific). Tissue sections were collected on glass slides (Fisherbrand™ Superfrost plus, Pittsburgh, PA), and stored at room temperature.

Prior to histological staining, tissue sections were deparaffinized and rehydrated in PBS. Staining kits for Hematoxylin (GHS280, Sigma Aldrich) & Eosin Y (HT1101128, Sigma Aldrich) solutions (H&E) and Masson’s trichrome stain (HT15-1KT, Sigma Aldrich) were used to assess cellularity and capsule thickness, respectively. For immunofluorescence staining, sections were subjected to antigen retrieval using heated citrate buffer (ab93678, Abcam), and incubated in blocking solution prepared using 4% normal goat serum, 1% bovine serum albumin, Fc block (anti-mouse CD16/32, 1:100, BD Biosciences), and 0.05% Triton-X in 1X PBS for 45 mins. Sections were then incubated in primary antibodies overnight at 4 °C. Primary antibodies used in this study include mouse anti-αSMA (1:500, Abcam, cat# ab7817), rabbit anti-Iba1(1:400, Wako, cat# 019-19741), and rabbit anti-myeloperoxidase (1:400, Agilent Technologies, cat# A0398). After three PBS washes, sections were incubated in goat anti-mouse IgG Alexa Fluor 568 (1:400, Abcam) and goat anti-rabbit IgG Alexa Fluor (1:400, Abcam) for 60 mins at room temperature. 4’,6-diamidino-2-phenylindole hydrochloride (DAPI) was used to counterstain nuclei. Sections were washed in PBS and anti-fade mounting media (Vectashield product H-1400, Vector Labs) was applied. Slides were cover slipped and samples were stored protected from light at 4°C until images were acquired.

### 2.5 Imaging and image analysis

Brightfield and fluorescence images were acquired with a Keyence microscope (BZ-X810) using Plan Fluor 20X (NA 0.45) and Plan Fluor 40X (NA .60) objective lenses. Nuclei count and capsule thickness measurements were carried out by an individual blinded to the study groups. H&E images (**Figure 1C**) acquired with a 40X objective were used to calculate cellularity. In ImageJ, a circle of radius 10 μm was generated as an ROI and was applied adjacent to the implant edge at 5-7 different locations per tissue section. Nuclei counts from multiple ROIs (per section) were averaged, normalized to total cell count obtained from a region adjacent to the implant site, and reported as fold change. 2-3 sections per animal were used. To measure capsule thickness, skin samples were stained with Masson’s trichrome and were imaged using a 20x objective. Stitched images were generated from tile scans and used in analysis (**Figure 2A**). The distance between the superficial side of the implant and the panniculus carnosus, the muscle layer below the hypodermis layer, was measured at 5 evenly spaced locations along the implant footprint (as outlined in **Figure 2B**) to obtain the average capsule thickness. 3-4 sections per animal were used to measure capsule thickness.

### 2.6 Statistical Analyses

All statistical analyses were carried out in JMP statistical software (version 14.0, SAS, Cary, NC). Statistical significance among groups were evaluated using a non-parametric, Kruskal-Wallis test with multiple comparisons using the Steel-Dwass method. Data presented in figures and text represent mean ± standard error of mean.

## Acknowledgements

This work was supported by funding from the National Institutes of Health R35GM128831 (C.S.S.) and National Science Foundation Graduate Student Fellowship under Grant No. DGE-1842473 (M.D.).

## Author Contributions

J.G., M.D., and C.S.S. conceived and designed the experiments. J.G., M.D., and V.S. carried out the experiments. J.G. and M.D. analyzed the results. C.S.S. secured the funding. J.G. wrote the first draft of the manuscript. All authors reviewed, provided feedback, and approved the manuscript.

## Data Availability

All data collected for this study are presented in the manuscript. Raw data will be available from the corresponding author on reasonable request.

